# Cell- and non-cell-autonomous ARF3 coordinates meristem proliferation and organ patterning in Arabidopsis

**DOI:** 10.1101/2022.01.12.476103

**Authors:** Ke Zhang, Hao Zhang, Yanyun Pan, Lin Guo, Shijun Tian, Jiarong Wei, Yunze Fu, Cong Wang, Ping Qu, Liantao Liu, Yongjiang Zhang, Hongchun Sun, Zhiying Bai, Jingao Dong, Cundong Li, Xigang Liu

## Abstract

In cell–cell communication, non-cell-autonomous transcription factors play vital roles in controlling plant stem cell fate. We previously reported that AUXIN RESPONSE FACTOR 3 (ARF3), a member of the ARF family with critical roles in floral meristem maintenance and determinacy, has a distinct accumulation pattern that differs from the expression domain of its encoding gene in the shoot apical meristem (SAM). However, the biological meaning of this difference is obscure. Here, we demonstrate that *ARF3* expression is mainly activated at the periphery of the SAM by auxin, where ARF3 cell-autonomously regulates the expression of meristem–organ boundary-specific genes, such as *CUP-SHAPED COTYLEDON1-3* (*CUC1-3*), *BLADE ON PETIOLE1-2* (*BOP1-2*) and *TARGETS UNDER ETTIN CONTROL3* (*TEC3*) to determine organ patterning. We also show that ARF3 is translocated into the organizing center, where it represses cytokinin activity and *WUSCHEL* expression to regulate meristem activity non-cell-autonomously. Therefore, ARF3 acts as a molecular link that mediates the interaction of auxin and cytokinin signaling in the SAM while coordinating the balance between meristem maintenance and organogenesis. Our findings reveal an ARF3-mediated coordination mechanism through cell–cell communication in dynamic SAM maintenance.

## INTRODUCTION

Multicellular organisms possess groups of cells with diverse identities and specific responsibilities. To orchestrate this diversity, complex signaling networks control stem cell fate based on hard-wired developmental programs (cell-autonomous effects) and environmental signals (non-cell-autonomous effects) (Gaillochet et al., 2017; Pfeiffer et al., 2017). Non-cell autonomy relies on two forms of intercellular communication: ligand-receptor-mediated perception that occurs in the apoplast and involves peptides or compounds such as phytohormones (Matsubayashi, 2003) and direct symplastic transport of signaling molecules such as mRNAs, siRNAs and non-cell-autonomous transcription factors (TFs) from cell to cell (Kurata et al., 2005).

In Arabidopsis (*Arabidopsis thaliana*), the shoot apical meristem (SAM) gives rise to all aerial structures over a plant’s lifetime (Miwa et al., 2009). Stem cells are located in the center zone (CZ) of the SAM, consisting of three cell layers. The CZ is surrounded by the peripheral zone (PZ), which is composed of transit amplifying cells derived from the stem cell niche and will differentiate into organ primordia with much higher cell division rates (Gaillochet and Lohmann, 2015). The organizing center (OC), located below the CZ, plays a vital role in stem cell maintenance (Carles and Fletcher, 2003). The SAM maintains the balance between stem cell self-renewal and indefinite lateral organ initiation and also establishes organ boundaries to properly sustain continuous growth (Lee et al., 2019; Zadnikova and Simon, 2014). After transition to reproductive development, the inflorescence meristem (IM) produces floral meristems (FMs) with a regular pattern of 137.5° between consecutive FMs (Chandler, 2012; Reinhardt et al., 2000). The FM contains stem cell populations during early growth stages to generate floral organs of genus- or species-specific size and number (Lee et al., 2019; Sun et al., 2009). In contrast to the SAM, each FM is genetically programmed to terminate stem cell activities after the initiation of carpels (Cao et al., 2015a; Chang et al., 2020; Sun and Ito, 2015; Sun et al., 2009).

The intertwined communication systems required for meristem maintenance, primordium initiation and organ boundary formation have been identified, including local transcriptional networks and non-cell-autonomous phytohormone signals (Brand et al., 2000; Cao et al., 2015b; Gaillochet et al., 2015; Gordon et al., 2009; Jasinski et al., 2005; Lee et al., 2019; Sun and Ito, 2015; Sun et al., 2009; Zadnikova and Simon, 2014). The *WUSCHEL* (*WUS*)–*CLAVATA* (*CLV*) feedback regulatory pathway plays a critical role in the maintenance of the stem cell pool. *WUS*, encoding a homeobox TF, is specifically expressed in the OC to specify stem cell fate by activating *CLV3* expression in the CZ in a non-cell-autonomous manner (Daum et al., 2014; Yadav et al., 2011). *CLV3* encodes a signal peptide that in turn represses *WUS* expression by binding to the CLV1 receptor complex. In addition, both cytokinins and auxin contribute to the fine-tuning of meristem maintenance in the SAM and FM (Lee et al., 2019). Cytokinins show a peak accumulation in the OC and control cell division by promoting *WUS* expression in the meristem (Chickarmane et al., 2012; Gordon et al., 2009; Riou-Khamlichi et al., 1999; Schaller et al., 2015). Sites of primordia formation are characterized by auxin maxima, while auxin accumulates to low levels in the OC of the SAM to determine organ patterning (Schaller et al., 2015; Shi et al., 2018; Vernoux et al., 2010). Auxin interacts both antagonistically and synergistically with cytokinins to regulate SAM and FM activity (Schaller et al., 2015). WUS restricts auxin signaling and response pathways in apical stem cells, where a low level of auxin signaling output is required for stem cell maintenance (Ma et al., 2019). However, the interactions between auxin and cytokinins within and between meristem zones as well as with *WUS* in meristem maintenance remain elusive. Our previous findings showed that AUXIN RESPONSE FACTOR3 (ARF3, also named ETTIN [ETT]) promotes FM determinacy by repressing cytokinin biosynthesis and signaling (Zhang et al., 2018). ARF3 can directly inhibit the expression of *ISOPENTENYLTRANSFERASE* (*IPT*) family members, encoding cytokinin biosynthetic enzymes, and *ARABIDOPSIS HISTIDINE KINASE4* (*AHK4*), encoding a cytokinin receptor, to reduce cytokinin activity (Cheng et al., 2013; Zhang et al., 2018). How ARF3 mediates the interaction between auxin and cytokinins is unclear. The dynamic maintenance of meristems depends on a precise balance between meristem self-renewal and lateral organ formation. MONOPTEROS (MP, also named ARF5) acts downstream of auxin and plays a key role in primordium initiation (Lee et al., 2019; Zhao et al., 2010). ARF3, ARF4 and MP are essential for organogenesis. ARF3 directly, and MP indirectly, represses *SHOOT MERISTEMLESS* (*STM*) expression to control organogenesis (Chung et al., 2019). *STM*, encoding a mobile KNOTTED-like homeobox TF repressing cell differentiation, is expressed throughout the SAM but is downregulated in incipient organ primordia (Chang et al., 2020; Jasinski et al., 2005; Long et al., 1996; Scofield et al., 2018). Formation of successive lateral organs on the flanks of the SAM requires the establishment of boundaries between meristems and organs to separate these two cell groups. Establishment of organ boundaries is determined by auxin accumulation due to directional auxin transport (Reddy et al., 2004; Vernoux et al., 2010; Zhao et al., 2013). *CUP-SHAPED COTYLEDON1–3* (*CUC1–3*) regulate the specification of organ boundaries (Bilsborough et al., 2011; Peaucelle et al., 2007): auxin regulates the establishment of organ boundaries by regulating the expression of *CUC2* and *BLADE ON PETIOLE* (*BOP*) (Zhao et al., 2013). In Arabidopsis, the divergence angle between successive primordia is approximately 137.5°. Loss-of-function mutants of boundary genes often cause alterations in phyllotactic patterning, such as the divergence angle, or internode length (Bencivenga et al., 2016; Peaucelle et al., 2007). Plants lacking ARF3 activity display dramatically altered phyllotaxis, as ARF3 regulates many target genes in an auxin-dependent manner, such as *TARGETS UNDER ETTIN CONTROL* (*TEC*): *TEC1* (corresponding to the TF gene basic helix-loop-helix [bHLH] *bHLH094*), *TEC2*, and *TEC3* (Byrne et al., 2003; Simonini et al., 2017). However, *arf3* also shows defects in floral patterning, such as more sepals and petals and fewer stamens (Nemhauser et al., 2000; Sessions et al., 1997). How ARF3 regulates pattern formation is unclear.

ARF3 plays a crucial role in regulating FM determinacy, gynoecium morphogenesis, self-incompatibility, phyllotactic patterning and floral organ patterns during floral development (Nemhauser et al., 2000; Sessions et al., 1997; Tantikanjana and Nasrallah, 2012; Zhang et al., 2018). *In situ* hybridization analysis revealed that *ARF3* is transcribed in clusters of cells giving rise to new FMs and floral organ primordia, which are similar to auxin maxima in these regions (Sessions et al., 1997). However, ARF3 is distributed throughout the IM and FM at stages 1–2, suggesting that ARF3 may coordinate meristem development in a non-cell-autonomous manner (Liu et al., 2014). In this study, we demonstrate that ARF3 migrates from the PZ into adjacent cells of the OC after being induced by auxin. ARF3 therefore controls organ patterning cell autonomously and regulates meristem activity non-cell-autonomously by mediating the interaction between auxin and cytokinins.

## RESULTS

### ARF3 regulates meristem activity and organ patterning

To investigate the roles of ARF3 in regulating meristem maintenance, we characterized the meristem phenotypes of the *arf3-29* mutant (Liu et al., 2014). We previously demonstrated that the SAM is larger in the *arf3* mutant than in its wild type L*er* (Figure 1A, 1B and 1K) (Zhang et al., 2018). *arf3-29* produced more L1 layer cells (the outermost layer of the SAM) (26.5 ± 2.9, n = 15) than L*er* (22.3 ± 1.8, n = 15) (Figure 1C, 1D and 1L), which was consistent with a previous report (Zhang et al., 2018). Moreover, the *arf3-29* mutant bore more siliques (62 ± 5.4, n = 21) than L*er* (38 ± 2.7, n = 25) and had a longer flowering period due to delayed global proliferative arrest (GPA) (Figure 1G, 1H, 1M and Supplemental Figure 1). We previously isolated *arf3-29* as an enhancer of FM determinacy defects in the weak *agamous* (*ag*) mutant allele *ag*-*10* (Liu et al., 2014), causing altered carpel polarity and enhanced FM indeterminacy in *ag-10 arf3-29* relative to *ag10*, with additional tissues in carpels (Figure 1E and 1F). These results indicated that ARF3 represses SAM activity by regulating meristem proliferation and stem cell termination.

**Figure 1.**
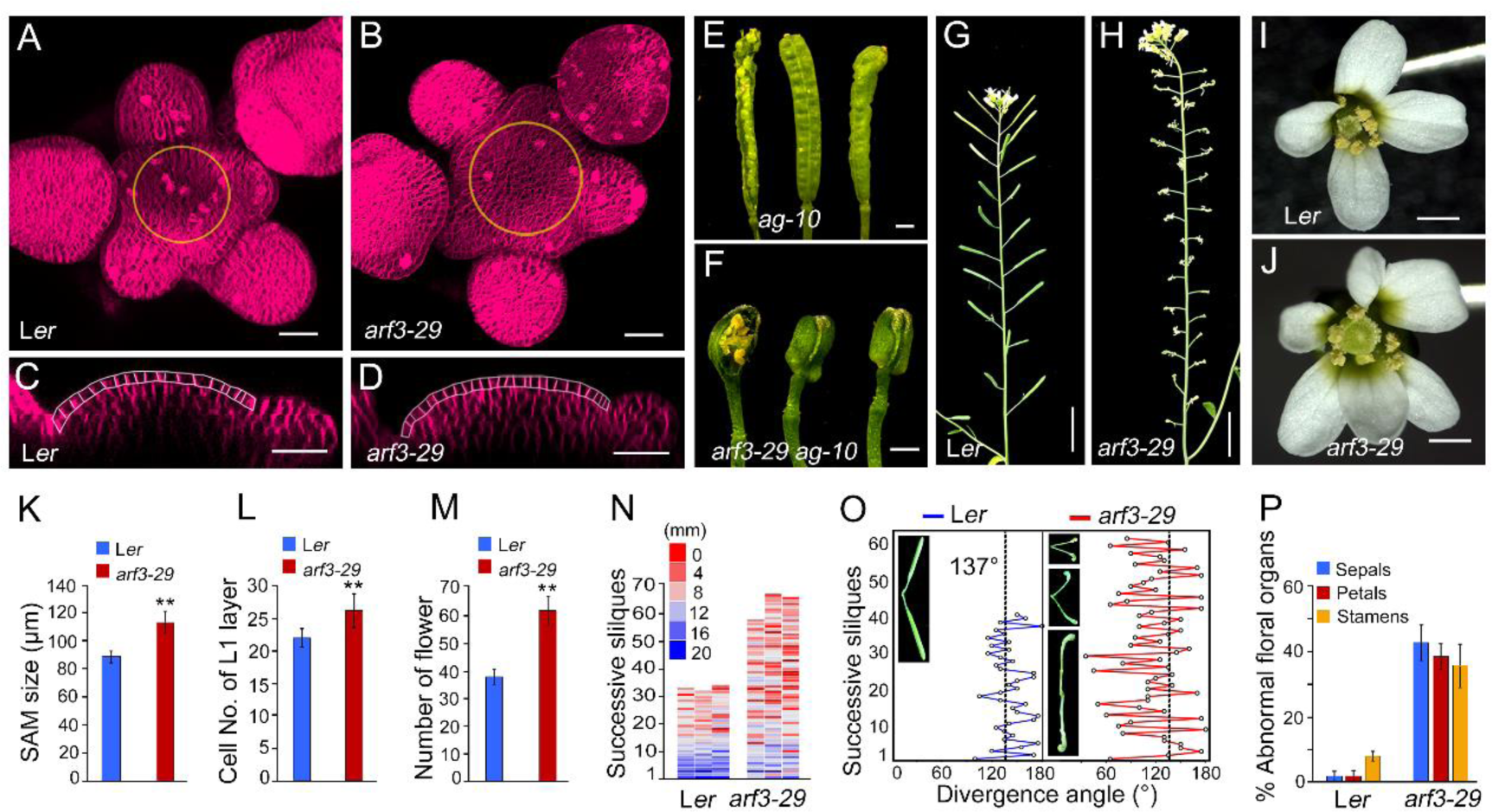
ARF3 regulates meristem activity and organ patterning. **A and B**, Representative 3D reconstructed top view of the SAM of L*er* (A) and the *arf3-29* mutant (B). **C and D**, Longitudinal section of the SAM in L*er* (C) and the *arf3-29* mutant (D). **E and F**, The *arf3-29* mutant enhances the FM determinacy defects of *ag-10*. Siliques in *ag-10* plants (E) and *ag-10 arf3-29* plants (F). **G and H**, Representative siliques on the main stem of L*er* (G) and the *arf3-29* mutant (H) 14 days after bolting. **I and J**, Flowers of wild type (L*er*) (I) and *arf3-29* (J). **K**, Size of the SAM in L*er* (n = 16) and *arf3-29* (n = 16) 7 days after bolting. \*\**P* < 0.01 (Student’s t test). **L**, Cell number of the SAM L1 layer in L*er* (n = 15) and *arf3-29* (n = 15). \*\**P* < 0.01 (Student’s t test). **M**, Number of flowers on the main stem during one life cycle in L*er* (n = 25) and *arf3-29* (n = 21). \*\**P* < 0.01 (Student’s t test). **N**, Distribution of internode length between two successive flowers along the stem in representative L*er* and *arf3-29* plants. **O**, Divergence angles between successive siliques in representative L*er* and *arf3-29* siliques. The insets show a representative example for L*er* and *arf3-29*. **P**, Percentages of abnormal floral organs in L*er* (163 flowers, 5 plants) and *arf3-29* (327 flowers, 6 plants). Bars = 25 μm in A–D; 1 mm in E, F, I and J; 1 cm in G and H.

Previous studies indicated that *ARF3* participates in the correct emergence of reproductive organ primordia (Sessions et al., 1997; Simonini et al., 2017). The *arf3-29* mutant displayed dramatically altered phyllotaxis and floral organ patterning compared to the wild type (Figure 1G, 1J, 1N and 1P). The *arf3-29* inflorescence was compact with very short internodes (Figure 1N). The divergence angle was near 137.5° in L*er* (141.7°), while it was around 119.3° in *arf3-29* (Figure 1O). Moreover, *arf3-29* produced more floral organs (sepals: 4.4 ± 0.05; petals: 4.4 ± 0.06; stamens: 5.6 ± 0.11) than did L*er* (Figure 1I, 1J, 1P and Supplemental Table 1). These results showed that *ARF3* functions on the regular organ patterning.

### Dynamic patterns of *ARF3* mRNA and ARF3 protein

To investigate *ARF3* expression in meristems, we analyzed auxin activity during early floral development by reanalyzing published data dissecting the architecture of gene regulatory networks controlling flower development using *ap1 cal 35Spro:AP1-GR* plants accumulating a fusion protein between APETALA1 (AP1) and the rat glucocorticoid receptor (GR) (Chen et al., 2018). AP1-GR induces FM initiation and floral development upon treatment with dexamethasone (DEX) (Wellmer et al., 2006). The expression of several genes was induced shortly after FM initiation in the published dataset (Figure 2A and Supplemental Data 1) and we validated these results by quantitative reverse transcription PCR (RT-qPCR) (Figure 2B–2F). These included auxin transporter genes such as *PIN-FORMED1* (*PIN1*) and *AUXIN1* (*AUX1*), auxin signaling components like *ARF3, ARF4* and *ARF17*, and auxin response genes such as *Small Auxin-up RNA8/53/54* (*SAUR8/53/54*). These results indicated increased auxin activity during early floral development, consistent with the role of auxin in promoting floral organ initiation. In agreement, we determined that *ARF3* expression was repressed following treatment with the auxin biosynthesis inhibitor yucasin or with the auxin transport inhibitor N-1-naphthylphthalamic acid (NPA) but was induced by indole-3-acetic acid (IAA) treatment, in line with our previous findings (Figure 2G) (Zhang et al., 2018).

**Figure 2.**
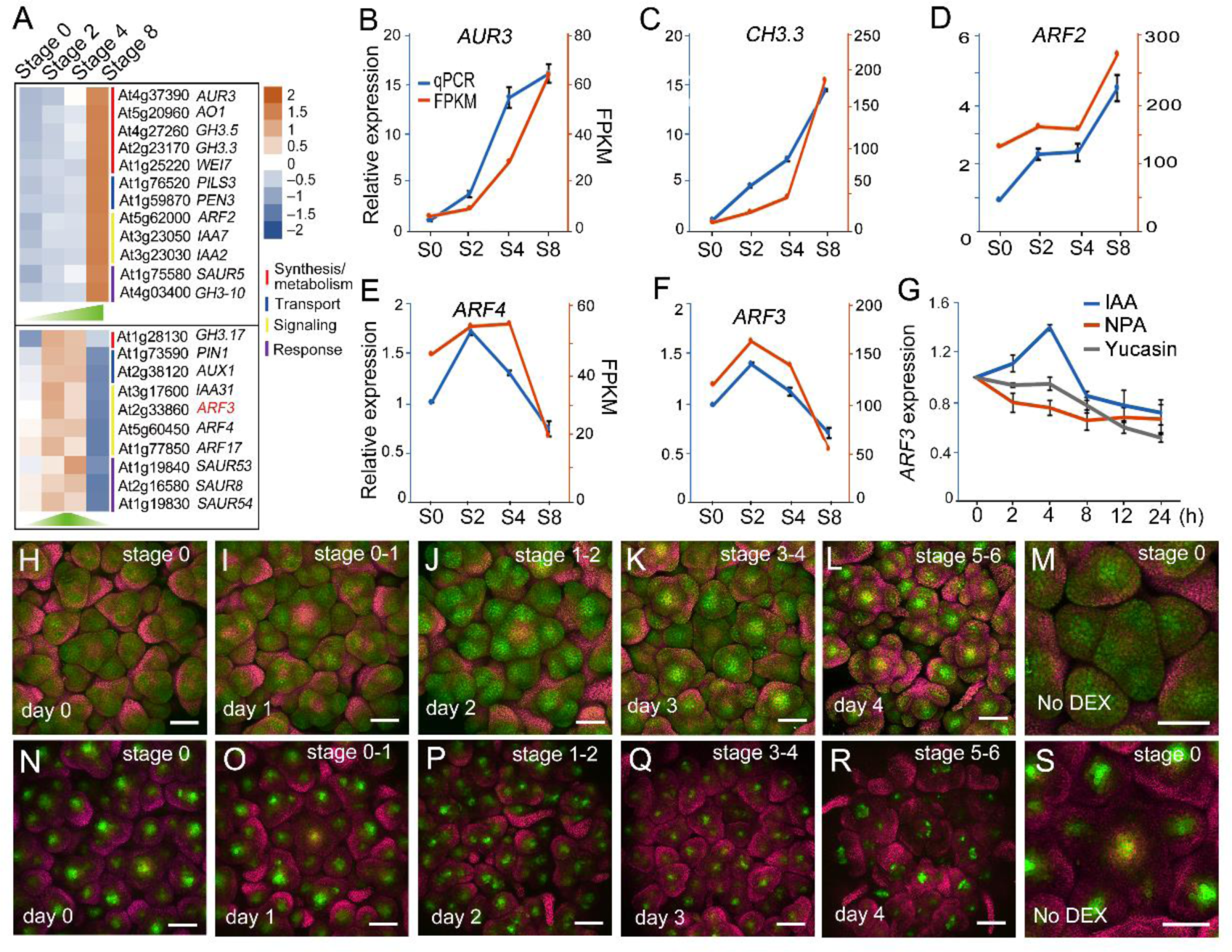
Dynamic patterns of *ARF3* expression and ARF3 distribution. **A**, Heatmap representation of differentially expressed gene expression patterns in auxin signaling during early floral development in *ap1 cal 35Spro:AP1-GR* (data from Chen et al., 2018). **B–F**, Comparison of expression levels of auxin-related genes estimated from published RNA-seq data (orange) and validated by RT-qPCR (blue). FPKM: fragments per kilobase of transcript per million mapped reads. qPCR: transcript levels measured by real-time RT-PCR. **G**, *ARF3* expression levels in L*er* inflorescences treated with IAA, NPA or yucasin. **H–M**, ARF3 distribution pattern during floral development in Arabidopsis. Comparison of ARF3-GFP protein abundance in *ap1 cal 35Spro:AP1-GR ARF3pro:ARF3-GFP* on day 0 (H), day 1 (I), day 2 (J), day 3 (K), day 4 (L) and No DEX (M) after a one-time treatment with 1 μM DEX. **N–S**, *TCSn:GFP* expression pattern during floral development in Arabidopsis. Comparison of GFP signals in *ap1 cal 35Spro:AP1-GR ARF3pro:ARF3-GFP* on day 0 (N), day 1 (O), day 2 (P), day 3 (Q), day 4 (R) and No DEX (S) after a one time treatment with 1 μM DEX. Bars = 100 μm in H–S.

To determine the distribution of ARF3 in meristems, we introduced *ARF3p:ARF3-GFP* encoding a functional fusion protein between ARF3 and the green fluorescent protein (GFP) into the *ap1 cal 35Spro:AP1-GR* background (Guo et al., 2018). After DEX treatment, ARF3 accumulation increased after FM initiation and peaked at stages 1-2 of early floral development with an even distribution throughout the meristem before becoming more concentrated in the OC at later floral development stages (Figure 2H-2L), illustrating the dynamic distribution pattern of ARF3. Given that auxin antagonistically interacts with cytokinins to regulate SAM and FM activity (Schaller et al., 2015), we also examined cytokinin signaling activity in the SAM by introducing the cytokinin reporter construct *TCSn:GFP* (Zürcher et al., 2013) into the *ap1 cal 35Spro:AP1-GR* background. Without DEX treatment, we observed an even distribution for cytokinin activity during very early stages (stages 0-1) of FM development (Figure 2N, 2O and 2S), in line with the role of cytokinins in maintaining stem cell activity by suppressing cell differentiation (Bartrina et al., 2011; Gordon et al., 2009; Sablowski, 2009). Following DEX treatment, cytokinin activity decreased and became restricted to the OC during late floral development (Figure 2N-2R). As ARF3 represses cytokinin biosynthesis and signaling (Zhang et al., 2018) and has a dynamic distribution (Figure 2H-2L), we hypothesized that ARF3 may mediate the dynamic repression of cytokinin activity by auxin during early floral development.

### ARF3 migrates from the PZ and CZ to the OC/niche

We previously showed that *ARF3* is expressed in clusters of cells that give rise to new FMs and floral organ primordia (Supplemental Figure 2A), while we detected ARF3 throughout the IM and stage 1-2 FMs (Figure 2M) (Liu et al., 2014), suggesting that ARF3 may act as a non-cell-autonomous TF. To confirm this hypothesis, we generated *ARF3pro:ARF3-nls-GFP* transgenic lines, harboring a transgene consisting of *GFP* with an in-frame nuclear localization signal (nls) downstream of *ARF3*. The addition of the nls restricted ARF3-nls-GFP to the nucleus, whereas ARF3-GFP localized to the cytoplasm, suggesting that the nls is effective in sequestering ARF3 to the nucleus and may prevent its cell-to-cell movement (Supplemental Figure 3). Longitudinal sections based on three-dimensional 3D reconstructions of IMs showed that ARF3-GFP signals are detected throughout the IM structure, including the CZ, PZ and OC zones containing the center region of L4-L6 layer cells (Figure 3A and 3B). However, we detected ARF3-nls-GFP signals only in L1-L3 layer cells but not in the OC zone (Figure 3E and 3F). In addition, we separately introduced *WUSpro:DsRed* (Liu et al., 2014), a *WUS* reporter specifically marking OC regions, into the *ARF3pro:ARF3-GFP* and *ARF3pro:ARF3-nls-GFP* transgenic lines via crossing. Compared to the even distribution of ARF3-GFP in the IM and early FMs (Figure 3C), ARF3-nls-GFP signals appeared limited to the new FM primordia and showed a weaker signal in the PZ and CZ zones (Figure 3G). Moreover, DsRed thoroughly overlapped with ARF3-GFP in the OC but only partially overlapped with ARF3-nls-GFP in the cells under the L2 layer (Figure 3D and 3H). DsRed signals were much weaker in the *ARF3pro:ARF3-GFP* transgenic lines than in *ARF3pro:ARF3-nls-GFP* lines (Figure 3C, 3D, 3G and 3H), indicating that ARF3 represses *WUS* expression directly.

**Figure 3.**
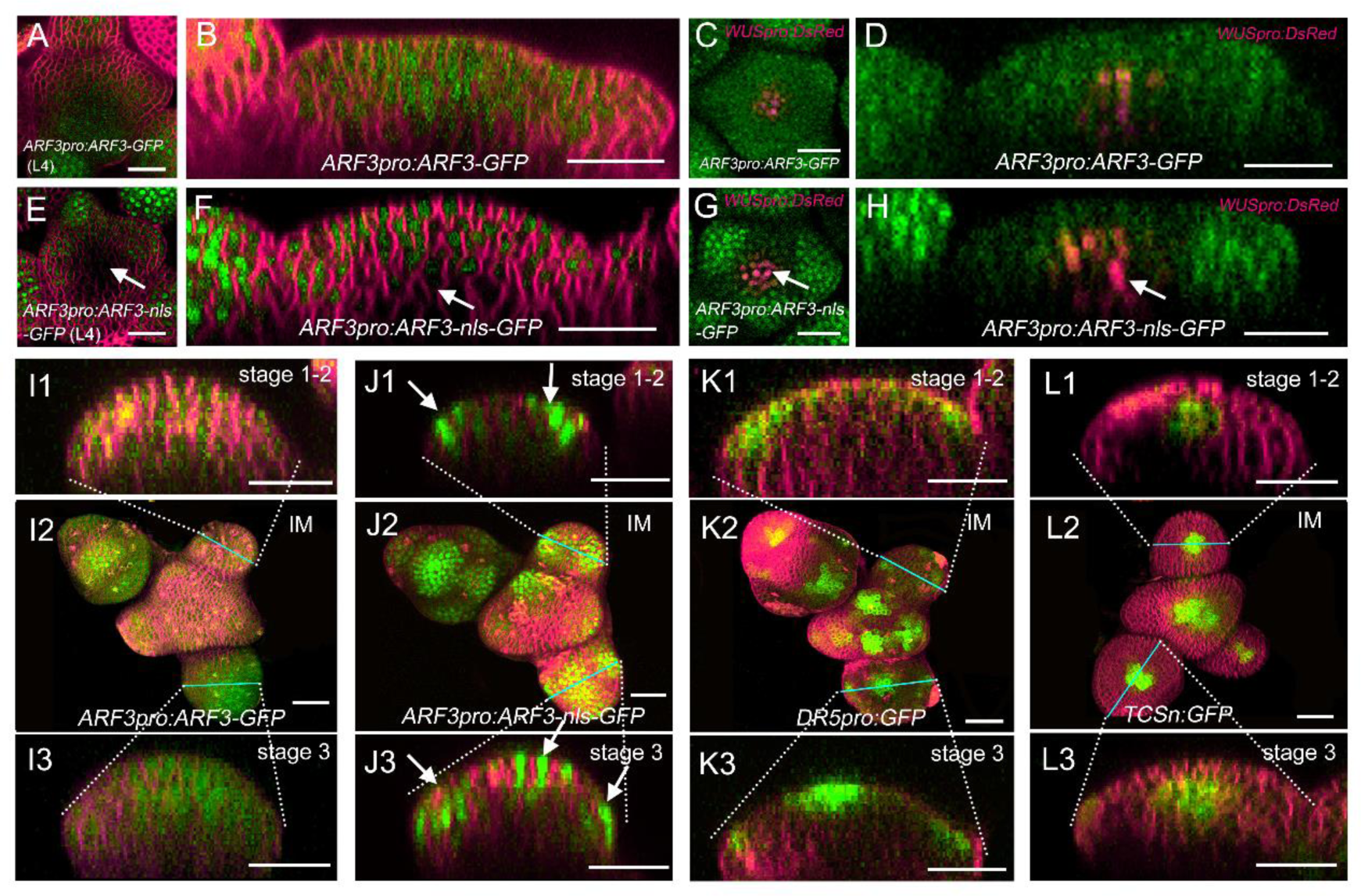
ARF3 protein migrates from cell to cell in the meristem. **A, B, E** and **F**, Transverse section and longitudinal section showing the distribution pattern of ARF3-GFP (A, B) and ARF3-nls-GFP (E, F) in inflorescence meristems. **C, D, G** and **H**, Transverse section and longitudinal section showing the distribution pattern of ARF3-GFP (C, D) and ARF3-nls-GFP (G, H) in *WUSpro:DsRed* (shown in magenta). **I1**–**I3**, ARF3-GFP distribution pattern in flowers. I1, early stages 1–2; I2, whole inflorescences and flower buds; I3, stages 2–3. **J1**–**J3**, ARF3-nls-GFP distribution pattern in flowers. J1, early stages 1–2; J2, whole inflorescences and flower buds; J3, stages 2–3. **K1**–**K3**, *DR5pro:GFP* expression in flowers. K1, early stages 1– 2; K2, whole inflorescences and flower buds; K3, stages 2–3. **L1**–**L3**, *TCSn:GFP* expression in flowers. L1, early stages 1–2; L2, whole inflorescences and flower buds; L3, stages 2–3. Bars = 25 μm.

To precisely explore the dynamic distribution of ARF3 in meristems, we examined the ARF3-GFP fluorescence signals in FMs at different stages. According to the 3D reconstructions, ARF3-GFP was ubiquitously distributed throughout IMs and early stages (1-3) of FMs in *ARF3pro:ARF3-GFP* lines (Figure 3I1-3I3). By contrast, ARF3-nls-GFP signals were enriched at the initiation sites of new organ primordia and the CZ (Figure 3J1–3J3), which also displayed high auxin activity, as evidenced by the synthetic auxin response reporter *DR5:GFP* (Figure 3K1-3K3), consistent with the induction of *ARF3* expression by auxin (Figure 2G) (Cheng et al., 2013). The *TCSn:GFP* reporter established that cytokinin activity is restricted to the OC during very early FM development (Figure 3L1-3L3). These findings indicated that auxin may induce *ARF3* expression, after which ARF3 migrates from the PZ and CZ to the OC/niche to regulate organ initiation or patterning during early floral development (stages 1-3).

### ARF3 migration is required for proper SAM activity maintenance

We further investigated the role of ARF3 movement in the regulation of SAM activity. The *arf3-29* mutant produced a larger SAM than L*er*, in agreement with enhanced SAM activity (Figure 1A, 1D and Figure 4A, 4B and 4E). The *arf3-29 ARF3pro:ARF3-GFP* line displayed a normal SAM size, indicating that ARF3-GFP fully rescues the *arf3-29* mutant phenotype (Figure 4C and 4E). We also crossed *ARF3pro:ARF3-nls-GFP* to *arf3-29* to assess whether ARF3-nls-GFP, unable to migrate from cell to cell, would function as ARF3-GFP. The *arf3-29 ARF3pro:ARF3-nls-GFP* line had SAMs intermediate in size between *arf3-29* and L*er* (Figure 4D and 4E). SAM size was positively correlated with the number of cells in the L1 layer across the four genotypes tested here (Supplemental Figure 4). These results suggested that the migration of ARF3 protein is required for SAM activity maintenance. Extended SAM activity resulted in delayed GPA responsible for the rise in flower number in the *arf3-29* mutant and in the *arf3-29 ARF3pro:ARF3-nls-GFP* line (Figure 4F and Figure 5A).

**Figure 4.**
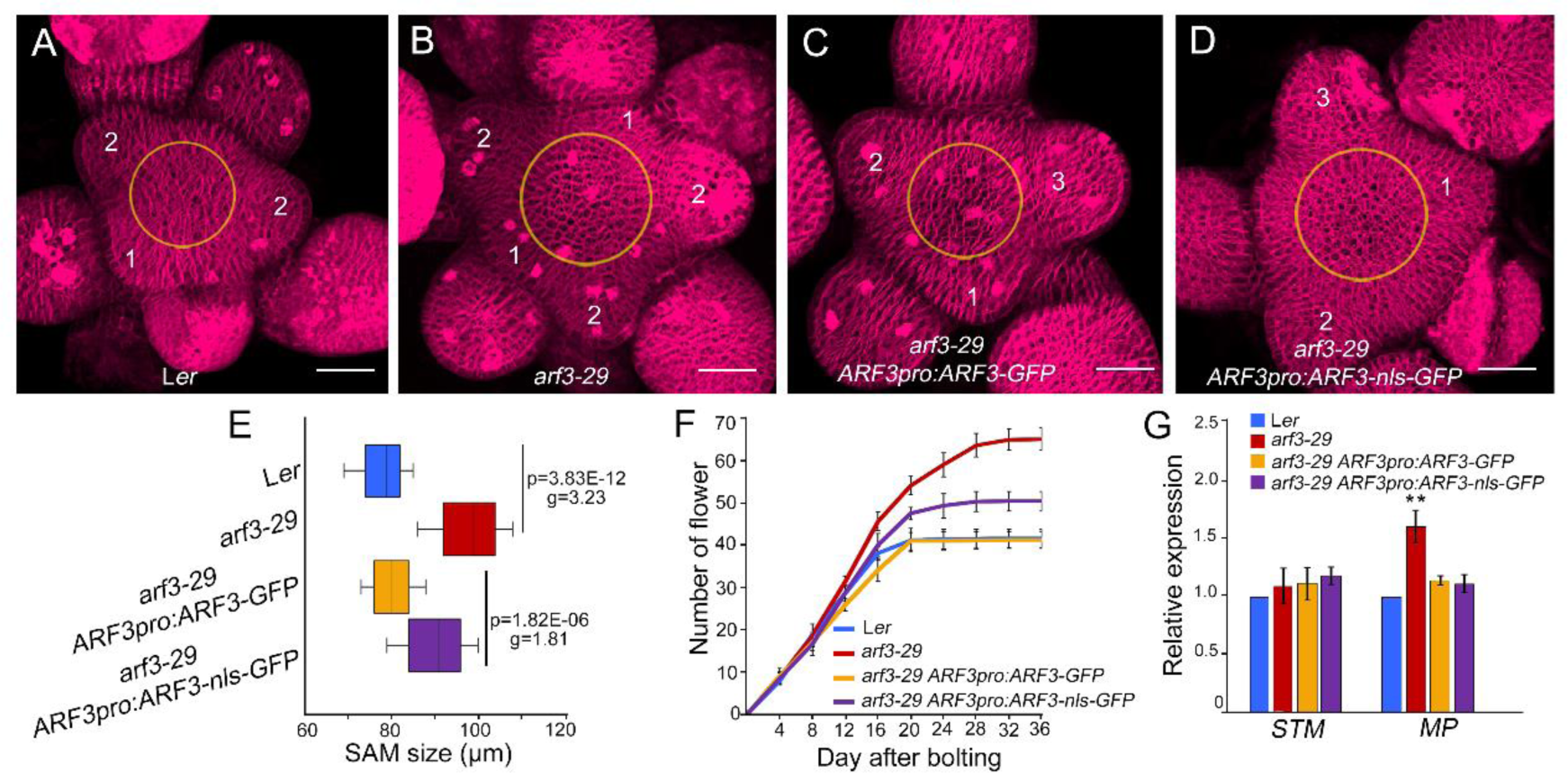
ARF3 independently regulates meristem maintenance and organ patterning. **A–D**, Representative top view of SAM size in the indicated genotypes. **E**, Boxplot representation of SAM size distribution in the indicated genotypes. **F**, Cumulative flower number over time in L*er* (n = 15), *arf3-29* (n = 15), *arf3-29 ARF3pro:ARF3-GFP* (n = 15) and *arf3-29 ARF3pro:ARF3-nls-GFP* (n = 15). **G**, Relative *STM* and *MP* expression levels in the indicated genotypes. \**P* < 0.05 (Student’s t test). Bars = 25 μm in A**–**D.

**Figure 5.**
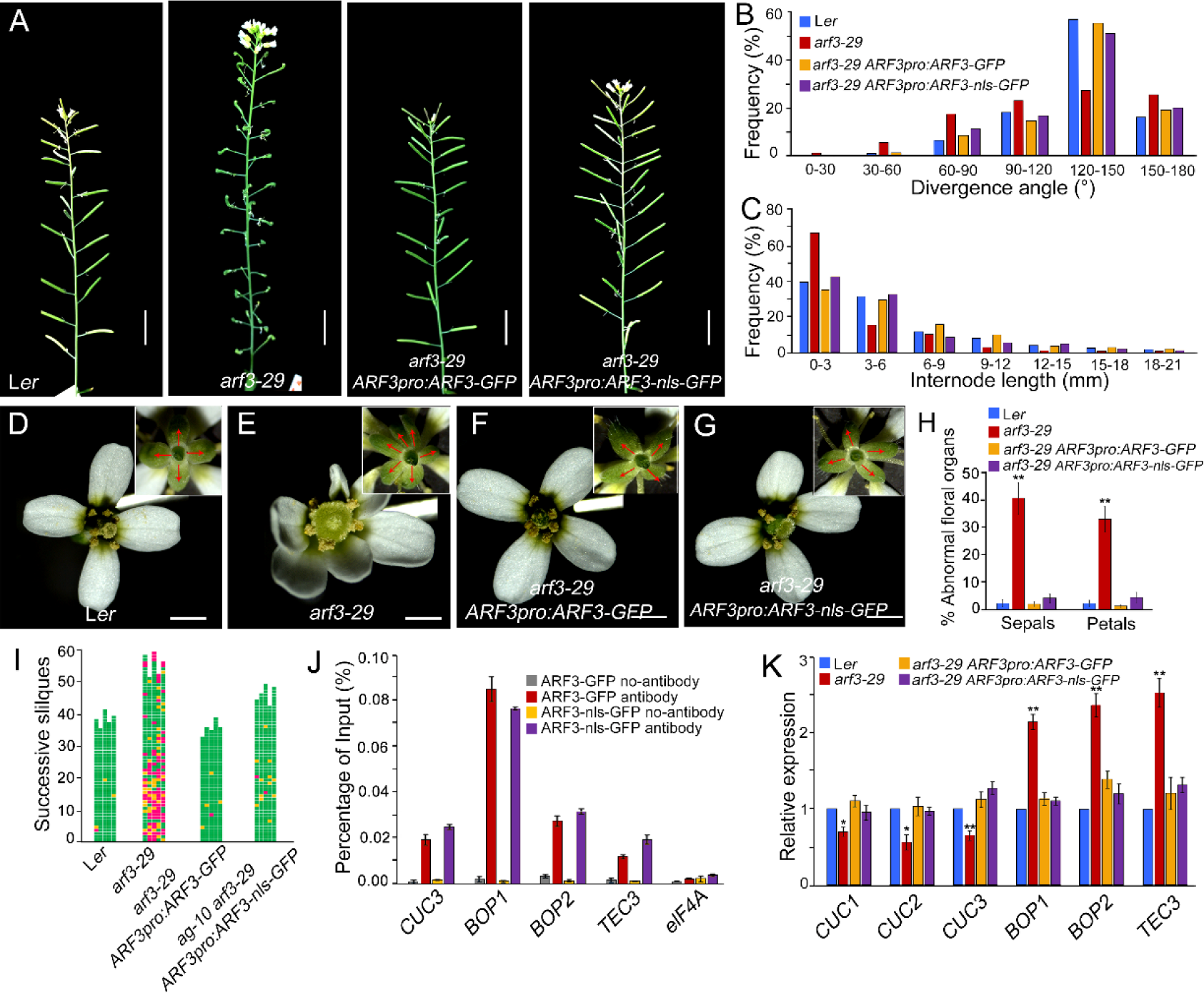
ARF3 controls phyllotactic pattern cell-autonomously. **A**, Representative images of inflorescence stems for the indicated genotypes. Scale bars = 1 cm. **B**, Distribution of divergence angle between two successive flowers, in 30° intervals. **C**, Distribution of internode length between two successive flowers, in 3-mm intervals. **D–G**, Flowers and floral organs for the indicated genotypes. Insets show the sepals marked by red arrows. **H**, Percentages of abnormal floral organs in L*er* (173 flowers from 4 plants), *arf3-29* (273 flowers from 5 plants), *arf3-29 ARF3pro:ARF3-GFP* (183 flowers from 5 plants) and *arf3-29 ARF3pro:ARF3-nls-GFP* (184 flowers from 4 plants). \*\**P* < 0.01 (Student’s t test). **I**, Schematic diagrams of floral organs formation in the indicated genotypes. Each column represents a single plant, and each square represents a flower. Green, normal flower with four sepals and four petals; yellow, flower with abnormal number of sepals or petals; magenta, flower with abnormal number of sepals and petals. **J**, ChIP assay for ARF3-GFP binding with anti-GFP antibody at *CUC3P1, BO2P1* and *TEC3* in *arf3-29 ARF3pro:ARF3-GFP* and *arf3-29 ARF3pro:ARF3-nls-GFP* inflorescences. The fragments examined are shown in Supplemental Figure 6. *eIF4A* (*EUKARYOTIC TRANSLATION INITIATION FACTOR4A*) served as a negative control. Error bars represent standard deviation (SD) from three biological repeats. **K**, Relative *CUC1–3, BOP2* and *TEC3* transcript levels in the indicated genotypes. \**P* < 0.05 and \*\**P* < 0.01 (Student’s t test). Bars = 1 cm in A, 1 mm in D–G.

The *ARF3pro:ARF3-nls-GFP* transgene fully rescued the organ initiation defect of *arf3-29* (compare Figure 4D to Figure 4A and 4B), indicating that proper organ initiation does not require ARF3 migration. ARF3, ARF4 and MP (ARF5) promote flower organogenesis by repressing *STM* expression (Chung et al., 2019), prompting us to measure *MP* and *STM* expression by RT-qPCR. While *STM*, the target of MP and ARF3, was expressed to comparable levels in L*er, arf3-29 ARF3pro:ARF3-GFP* and *arf3-29ARF3pro:ARF3-nls-GFP, MP* expression increased significantly in *arf3-29*, which was rescued by both *arf3-29 ARF3pro:ARF3-GFP* and *arf3-29 ARF3pro:ARF3-nls-GFP* transgenes (Figure 4H). We concluded that ARF3 migration in the SAM is required for the maintenance of meristem activity but not for organ initiation.

### ARF3 controls organ patterning cell-autonomously

To dissect the molecular mechanism underlying the cell-autonomous role of ARF3 in organ initiation, we compared the silique emergence sites and internodes in L*er, arf3-29, arf3-29 ARF3pro:ARF3-GFP* and *arf3-29 ARF3pro:ARF3-nls-GFP*, as they showed different phyllotaxy patterns (Figure 5A). The average divergence angle between successive primordia in *arf3-29* inflorescence apices was 123.3° ± 39.64 (n = 205), which was smaller than the theoretical angle of 137.5°. Only 27.3% of the divergence angles fell into the 120-150° range in the mutant (Figure 5B), compared to 56.5% in L*er*, 55.2% in *arf3-29 ARF3pro:ARF3-GFP* and 51.2% in *arf3-29 ARF3pro:ARF3-nls-GFP* (Figure 5A and 5B). We also measured internode length between successive organs along the main inflorescence stem. Most internodes ranged from 0 mm to 3 mm in length (67.9% of total tested plants) in *arf3-29* (Figure 5C), whereas the other genotypes displayed fewer internodes in this range: 40.0% for L*er*, 35.5% in *arf3-29 ARF3pro:ARF3-GFP*, and 43.4% in *arf3-29 ARF3pro:ARF3-nls-GFP*. Internodes tended to be longer in these genotypes, with 31.7% (L*er*), 29.9% (*arf3-29 ARF3pro:ARF3-GFP*), and 33.1% (*arf3-29 ARF3pro:ARF3-nls-GFP*) in the 3- to 6-mm range, compared to only 15.9% for *arf3-29* (Figure 5C). These results suggested that ARF3 controls the phyllotactic pattern in a cell**-**autonomous manner. The *arf3-29* mutant produced abnormal flowers with more sepals and petals but with fewer stamens, indicative of impaired floral organ initiation (Figure 5D, 5E, 5H and Supplemental Figure 5) (Sessions et al., 1997). Both *arf3-29 ARF3pro:ARF3-GFP* and *arf3-29 ARF3pro:ARF3-nls-GFP* lines exhibited the same number of floral organs as L*er* (Figure 5F-5I). Neither *ARF3pro:ARF3-GFP* nor *ARF3pro:ARF3-nls-GFP* fully rescued the aberrant number of stamens, suggesting that ARF3 may affect stamen development via some unknown mechanism(s) (Supplemental Figure 5).

*TEC3* participates in spiral phyllotaxis formation, as well as the meristem-organ boundary-specific genes, such as *CUC3, BOP1* and *BOP2*; all are target genes of ARF3 (Simonini et al., 2017). We performed chromatin immunoprecipitation followed by quantitative PCR (ChIP-qPCR) using stage 6 inflorescences and younger flowers to test whether ARF3 regulates these boundary-specific genes in a migration-dependent manner. We purified chromatin bound by ARF3-GFP or ARF3-nls-GFP with anti-GFP antibodies. Both ARF3-GFP and ARF3-nls-GFP bound to the *CUC3, BOP1, BOP2* and *TEC3* loci with similar enrichment levels in *arf3-29 ARF3pro:ARF3-GFP* and *arf3-29 ARF3pro:ARF3-nls-GFP* (Figure 5J and Supplemental Figure 6). Furthermore, *CUC1, CUC2* and *CUC3* transcript levels decreased, while the expression of *BOP1, BOP2* and *TEC3* increased, in the *arf3-29* mutant (Figure 5K). Introduction of the *ARF3pro:ARF3-GFP* and *ARF3pro:ARF3-nls-GFP* transgenes into the *arf3-29* mutant returned the expression of these genes to wild-type levels (Figure 5K). These findings suggest that ARF3 plays a role in phyllotactic patterning by regulating boundary-specific gene expression in a cell-autonomous manner.

### ARF3 controls SAM activity and FM determinacy non-cell-autonomously

Precise stem cell activity maintenance and termination are critical for SAM and FM maintenance and programmed FM determinacy (Chang et al., 2020). We previously demonstrated that ARF3 promotes FM determinacy by mediating the interaction between auxin and cytokinins (Zhang et al., 2018). We thus examined what role, if any, ARF3 migration may play in meristem maintenance and FM determinacy. Our working hypothesis was that ARF3 protein migration is required for FM determinacy. To test this idea, we separately introduced the *ARF3pro:ARF3-GFP* and *ARF3pro:ARF3-nls-GFP* transgenes into the *ag-10 arf3-29* double mutant, which displayed a severe FM determinacy defect, with unfused and bulging carpels with hyperplastic tissue and lacking seeds inside (Supplemental Figure 7B and 7F) (Liu et al., 2014; Zhang et al., 2018). The *ARF3pro:ARF3-nls-GFP* transgene only partially rescued the FM indeterminacy phenotype seen in *ag-10 arf3-29* (Supplemental Figure 7D and 7H), while *ARF3pro:ARF3-GFP* fully rescued this defect (Figure 7C and 7G). About 68.7% of *ag-10 arf3-29 ARF3pro:ARF3-nls-GFP* siliques appeared normal, as in *ag-10* or *ag-10 arf3-29 ARF3pro:ARF3-GFP*, but the remaining 31.3% of siliques exhibited bulging carpels with additional tissue growing inside and some seeds (Supplemental Figure 7I and 7J). These results indicated that ARF3 migration is required for proper FM determinacy.

**Figure 6.**
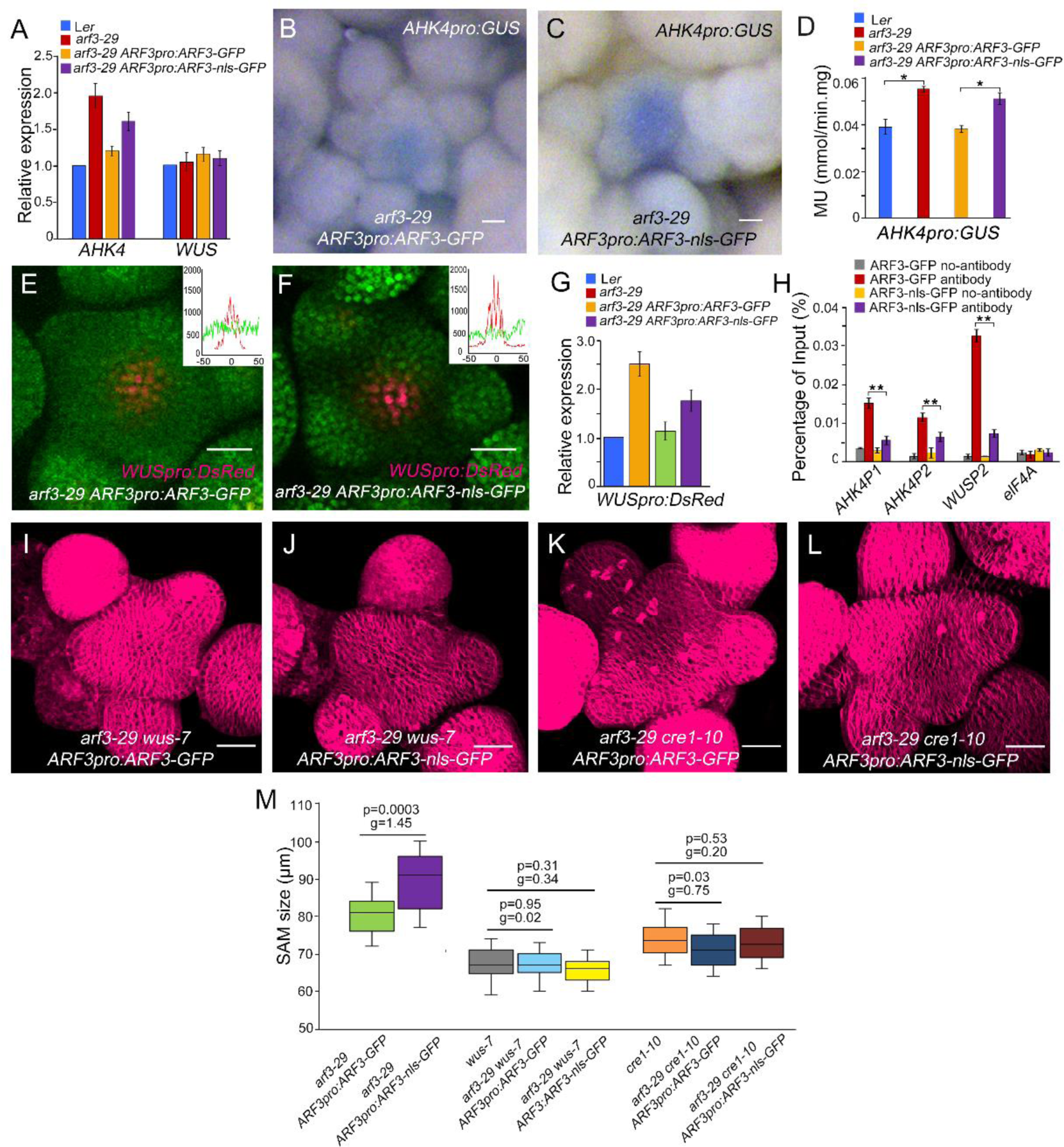
ARF3 controls meristem activity non-cell-autonomously. **A**, Relative *AHK4* and *WUS* transcript levels in the indicated genotypes. \**P* < 0.05 and \*\**P* < 0.01 (Student’s t test). **B and C**, *AHK4pro:GUS* expression pattern in *arf3-29 ARF3pro:ARF3-GFP* (B) and *arf3-29 ARF3pro:ARF3-nls-GFP* (C) inflorescences under the same staining conditions. **D**, Quantification of GUS activity in inflorescences of the indicated genotypes. Data are shown as means of three biological replicates with independently prepared inflorescence materials containing unopened flowers. \**P* < 0.05 (Student’s t test). **E and F**, *WUSpro:DsRed* expression (magenta) in the SAM of *arf3-29 ARF3pro:ARF3-GFP* (E) and *arf3-29 ARF3pro:ARF3-nls-GFP* (F). The insets show the average fluorescence signal of DsRed, ARF3-GFP and ARF3-nls-GFP signals. y-axis, signal intensity; x-axis, position along the SAM (μm). \**P* < 0.05 and \*\**P* < 0.01 (Student’s t test). **G**, Relative *DsRed* transcript levels in the indicated genotypes. **H**, ChIP assay with anti-GFP antibody to examine ARF3 binding to *AHK4P1, AHK4P2* and *WUSP2* in *arf3-29 ARF3pro:ARF3-GFP* and *arf3-29 ARF3pro:ARF3-nls-GFP* inflorescences. *eIF4A* served as a negative control. Error bars represent the SD from three biological repeats with independently prepared inflorescence materials containing unopened flowers. \*\**P* < 0.01 (Student’s t test) between *arf3-29 ARF3pro:ARF3-GFP* and *arf3-29 ARF3pro:ARF3-nls-GFP* inflorescences. **I–L**, Representative SAM top view in the indicated genotypes. **M**, Boxplot representation of SAM size distribution in the indicated genotypes. Statistical test: Student’ t test (p); Effect size: Hedges’ coefficient (g). Bars = 25 μm in B, C, E, F, I and L.

**Figure 7.**
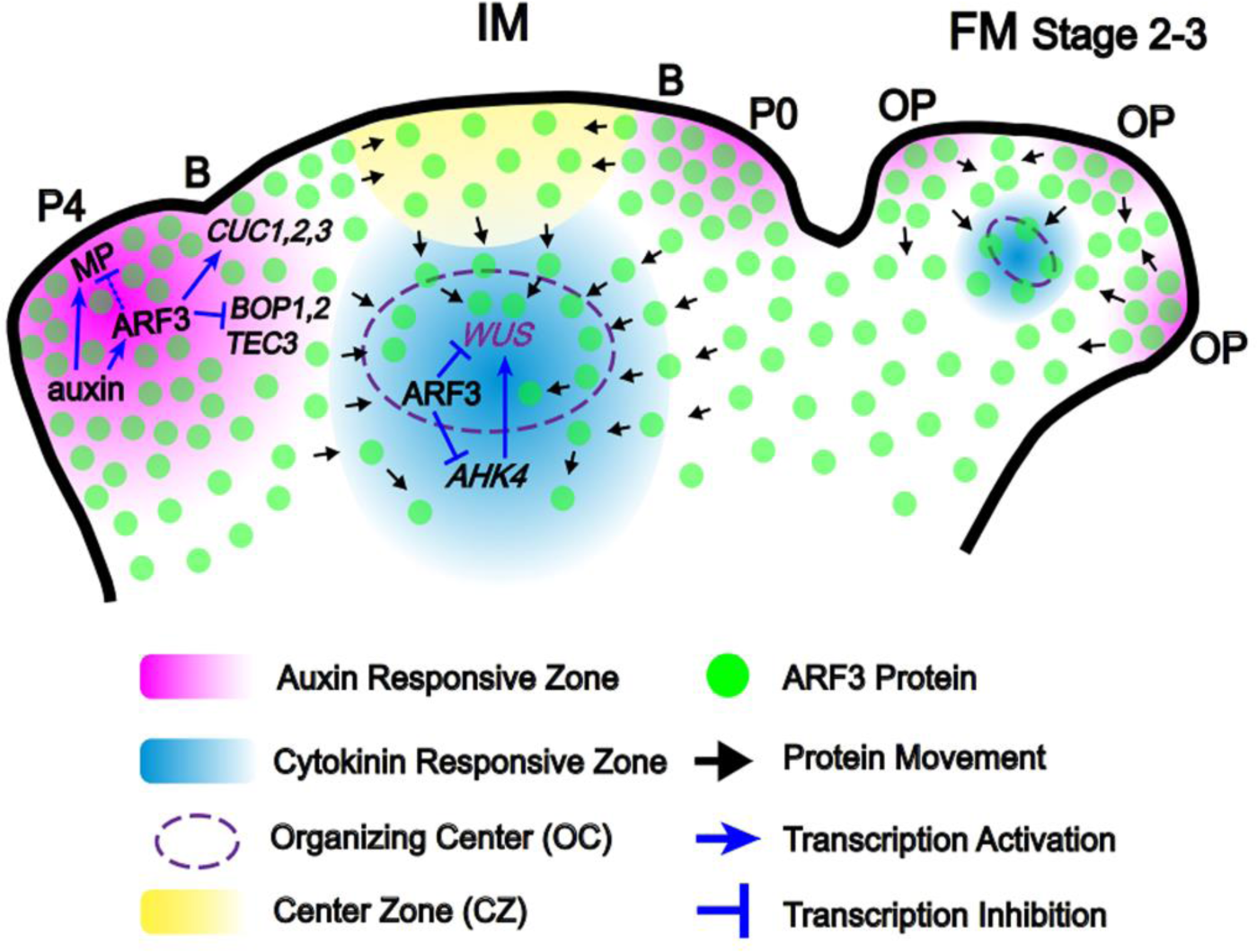
Model of meristem activity and phyllotactic pattern control by ARF3 in a non-cell- and cell-autonomous manner. In the IM, auxin promotes *ARF3* expression, which in turn regulates meristem–organ boundary-specific genes (*CUC1–3, BOP1–2* and *TEC3*) and *MP* in a cell-autonomous manner. ARF3 also migrates from the PZ to the OC, where it directly represses *AHK4* and *WUS* expression to control meristem activity in a non-cell-autonomous manner. P0 and P4, primordia of flower buds at different developmental stages; B, meristem–organ boundary; OP, flower organ primordia.

The WUS-CLV3 negative feedback regulatory loop is critical for stem cell maintenance and is fine-tuned by cytokinin signaling at the OC (Lee et al., 2019). We previously demonstrated that ARF3 promotes FM determinacy by repressing cytokinin signaling and biosynthesis (Liu et al., 2014; Zhang et al., 2018). To investigate the molecular mechanism underlying the role of mobile ARF3 in SAM maintenance and FM determinacy, we examined the expression of *WUS* and *ARABIDOPSIS HISTIDINE KINASE4* (*AHK4*), encoding a cytokinin receptor, in L*er, arf3-29, arf3-29 ARF3pro:ARF3-nls-GFP* and *arf3-29 ARF3pro:ARF3-nls-GFP* by RT-qPCR. While *WUS* expression was comparable in the inflorescences of all tested genotypes, *AHK4* expression was higher in *arf3-29* relative to L*er*, but returned to wild-type levels in *arf3-29 ARF3pro:ARF3-GFP*, and was intermediate between L*er* and *arf3-29* in *arf3-29 ARF3pro:ARF3-nls-GFP* (Figure 6A). We confirmed this observation qualitatively and quantitatively with the *AHK4pro:GUS* reporter line (Figure 6B-6D and Supplemental Figure 8), indicating that ARF3 migration is required for the repression of cytokinin activity by ARF3 in the OC. Since *WUS* is expressed in a very limited number of cells, we visualized the *WUS* expression pattern using a *WUSpro:DsRed* reporter construct. As evidenced by quantitative fluorescence analysis of DsRed signal and RT-qPCR detection of *DsRed, WUS* was derepressed in *arf3-29* and *arf3-29 ARF3pro:ARF3-nls-GFP* relative to L*er* and *arf3-29 ARF3pro:ARF3-GFP* (Figure 6E-6G). ChIP-PCR analysis revealed the stronger occupancy of ARF3 at the *WUS* and *AHK4* loci in *arf3-29 ARF3pro:ARF3-GFP* compared to *arf3-29 ARF3pro:ARF3-nls-GFP*.

The weak *wus* mutant allele *wus-7* and *ahk4* mutants have smaller SAMs (Supplemental Figure 9). We separately introduced *wus-7* and the *AHK4* mutant allele *cre1-10* (*cytokinin response1-10*) into *arf3-29 ARF3pro:ARF3-nls-GFP*, which revealed that both mutants fully suppress the enlarged SAM size phenotype (Figure 6I-6Q). These results demonstrated that ARF3 mobility is required for its non-cell-autonomous maintenance of SAM activity and FM determinacy by repressing cytokinin activity and *WUS* expression in the OC.

## DISCUSSION

SAM maintenance results from a balance between the rate of meristem proliferation and the rate of new organ primordium initiation, which is controlled by the WUS-CLV3 feedback regulatory loop and the interaction between STM and meristem–organ boundary genes, respectively (Chang et al., 2020). How these two mechanisms interact and which factors mediate the crosstalk between stem cell renewal and organ initiation are not clear. We demonstrated here that ARF3 independently represses organ initiation and stem cell renewal cell-autonomously and non-cell-autonomously, respectively (Figure 7). At the meristem–organ boundary and new organ primordium, ARF3 is translated locally and regulates the expression of *MP* and meristem-organ boundary genes such as *CUC1-3* and *BOP1-2* as well as *TEC3* to fine-tune organ initiation (Figure 5J-5K). In agreement, *ARF3* loss of function resulted in the initiation of ectopic FMs and floral organs, phenotypes that were rescued by an immobile version of ARF3 trapped in the nucleus (Figure 4A, 4B, 4D and Figure 5D, 5E and 5G). The *arf3 arf4 mp* triple mutant produces pin-like SAM structures, indicating that the combined activity of ARF3, ARF4, and MP is required for organ initiation (Chung et al., 2019). The differential rescue of the *arf3-29* mutant by ARF3-GFP and ARF3-nls-GFP indicates that ARF3 exerts a complex function in organ initiation, at the center of which the dynamic distribution of ARF3 or other interacting factors is essential. ARF3 migrating from the PZ to the OC directly repressed *WUS* expression (Figure 6E-H) (Liu et al., 2014), which regulates stem cell activity and leads to altered SAM GPA and FM determinacy in the *arf3-29* and *arf3-29 ag-10* mutants (Figure 5A and Supplemental Figure 7A, 7B, 7E and 7F). Nucleus-localized ARF3 failed to rescue the enlarged SAM of *arf3-29* or the FM indeterminacy of *arf3-29 ag-10* (Figure 4D-4E and Supplemental Figure 7D and 7H), demonstrating that ARF3 mobility is critical for its regulation of stem cell activity. Therefore, mobile ARF3 mediates the crosstalk between meristem differentiation and proliferation.

Auxin and cytokinins exhibit distinct activity patterns and antagonistically regulate SAM maintenance. Auxin maxima are detected in regions of primordia formation and meristem-organ boundaries, where auxin promotes meristem differentiation. By contrast, cytokinins are concentrated in the OC, where they regulate *WUS* expression to fine-tune stem cell proliferation (Schaller et al., 2015). Auxin induces *MP* expression, which in turn represses the expression of type-A *ARABIDOPSIS RESPONSE REGULATOR*s (*ARR*s) to regulate cytokinin activity in the OC. We determined that ARF3 mediates the repression of cytokinin activity and *WUS* expression by auxin (Zhang et al., 2018). WUS restricts and maintains minimal auxin activity in the CZ by globally regulating the auxin pathway (Ma et al., 2019), although the exact nature of the elaborate regulatory networks between auxin, cytokinins and *WUS* is still unclear. We showed here that auxin and cytokinins exhibit dynamic activity patterns during early FM development. Auxin showed an increased activity gradient in the developing FM, with high auxin activity concentrated at floral organ primordium initiation sites (Figure 3K1), consistent with its function in inducing cell differentiation. In addition, we detected strong auxin activity in the CZ of stage 3 FMs (Figure 3K3), which disappeared at later stages (Ma et al., 2019). By contrast, cytokinin activity was low during FM development and high in the OC (Figure 3L), where cytokinins promote *WUS* expression. ARF3 and MP mediate the interaction between auxin and cytokinin to regulate meristem maintenance (Zhang et al., 2018; Zhao et al., 2010). The *ARF3* expression pattern and the distribution pattern of immobile ARF3-nls-GFP highly overlapped with the auxin activity pattern in meristems (Figure 3J and Supplemental Figure 2), consistent with the induction of *ARF3* expression by auxin. We previously observed that ARF3 directly regulates cytokinins in the OC (Zhang et al., 2018). However, compared to ARF3-GFP, ARF3-nls-GFP failed to rescue the FM indeterminacy phenotype seen in *arf3-29 ag-10*, showing prolonged *WUS* expression and enhanced *AHK4* expression (Figure 6A–6G), indicating that cell-to-cell ARF3 mobility is required for its role in regulating stem cell activity. We thus unraveled an ARF3-mediated post-translational regulatory mechanism (Figure 7), whereby ARF3 is induced by auxin in the PZ and in new organ primordia before migrating to the OC, where it directly represses *WUS* expression and cytokinin activity to balance stem cell proliferation and meristem differentiation.

## METHODS

### Plant Materials and Growth Conditions

Mutants and transgenic *Arabidopsis thaliana* lines are in the L*er* background, except for *TCSn: GFP* (Zürcher et al., 2013), *ahk4* (originally *cre1-10*) (Higuchi et al., 2004), and *AHK4pro:GUS* (Higuchi et al., 2004), which are in the Col-0 background. *ag-10* (Ji et al., 2011), *arf3-29* (Liu et al., 2014), *arf3-29 ag-10* (Liu et al., 2014), *ARF3pro:ARF3-GFP* (Liu et al., 2014), *arf3 ag-10 ARF3pro:ARF3-GFP* (Zhang et al., 2018), *arf3 ARF3pro:ARF3-GR* (Zhang et al., 2018), *ap1 cal 35Spro:AP1-GR* (Liu et al., 2011), *WUSpro:DsRed* (Liu et al., 2014), and *wus-7* (Lin et al., 2016) were previously described. All plants were grown at 23°C under long-day conditions (16-h light [100 μmol m^−2^ s^−1^]/8-h dark).

### Vector Construction and Transformation

To construct *ARF3pro:ARF3-nls-GFP*, the *nls-GFP* fragment containing the sequence for the nuclear localization signal (*nls*) and *GFP* with stop codon was amplified by PCR using pMDC107 as a template (Liu et al., 2014) and then digested with AscI and inserted into the AscI restriction site of pMDC107*-ARF3pro:ARF3-GFP*. The resulting *ARF3pro:ARF3-nls-GFP* construct was transformed into *arf3-29* and *arf3-29 ag-10* via the floral dip method (Clough and Bent, 1998), and primary transformants were selected for resistance to hygromycin.

### Generation of Mutant Combinations

To produce *ap1 cal arf3 ARF3pro:ARF3-GFP 35Spro:AP1-GR* and *ap1 cal TCSn:GFP 35Spro:AP1-GR* lines, *arf3-29 ARF3pro:ARF3-GFP* and *TCSn:GFP* plants were crossed to *ap1 cal 35Spro:AP1-GR*. In the F_2_ population, *arf3-29* plants were identified by genotyping; *ap1 cal* plants were identified based on phenotypes; the presence of the *TCSn:GFP* transgene was determined by watering or spraying the plants with the herbicide; the *35Spro:AP1-GR* transgene was selected by growth on medium containing kanamycin; the *ARF3pro:ARF3-GFP* transgene was selected on medium containing hygromycin.

To produce *arf3 WUSpro:DsRed ARF3pro:ARF3s-GFP* and *arf3 WUSpro:DsRed ARF3pro:ARF3-nls-GFP* lines, *arf3 ARF3pro:ARF3-nls-GFP* and *arf3 ARF3pro:ARF3-GFP* were crossed to *WUSpro:DsRed*. Plants harboring the *WUSpro:DsRed* transgene were selected based on DsRed signals on a confocal microscope.

To produce the *arf3 AHK4pro:GUS ARF3pro:ARF3s-GFP* and *arf3 AHK4pro:GUS ARF3pro:ARF3-nls-GFP* lines, *arf3 ARF3pro:ARF3-nls-GFP* and *arf3 ARF3pro:ARF3-GFP* were crossed to *AHK4pro:GUS*. Plants carrying the *AHK4pro:GUS* transgene were selected based on positive GUS staining.

To produce the *arf3 cre1-10 ARF3pro:ARF3s-GFP, arf3 cre1-10 ARF3pro:ARF3-GFP, arf3 wus-7 ARF3pro:ARF3s-GFP, arf3 wus-7 ARF3pro:ARF3-nls-GFP* lines, *arf3 ARF3pro:ARF3-nls-GFP* and *arf3 ARF3pro:ARF3-GFP* were crossed to *cre-10* and *wus-7*. Plants homozygous for *cre1-10* or *wus-7* were selected by genotyping.

All primers used for genotyping are listed in Supplemental Data Set 1.

### Microscopy

Optical photographs were taken under a Leica M205d/ DFC450 stereoscopic microscope. Confocal images were taken with a Leica TCS SP8 confocal microscope according a previously described protocol (Wei et al., 2020). Appropriate filter sets and lasers were selected for fluorescence excitation and scanning. Chlorophyll autofluorescence was excited at 488 nm and detected in the 660- to 700-nm range. GFP was excited at 488 nm, and detected in the 510- to 550-nm range. For FM4-64, the excitation wavelength was 561 nm, and the detection range was 570-620 nm. All live imaging experiments were performed as previously described (Wei et al., 2020).

### Plant Treatments and Tissue Collection

For chemical treatments, inflorescences from 3-week-old plants were treated with 5 μM 6-BA (Sigma-Aldrich), 50 μM IAA (Sigma-Aldrich), 100 μM yucasin (Sigma-Aldrich), 1 μM DEX (Sigma-Aldrich) in DMSO along with 0.015% (v/v) Silwet L-77.

For RT-qPCR analysis, inflorescences were dissected under a stereomicroscope to remove stage 7 and older flowers; 20–40 flowers were pooled for RNA extraction for each biological replicate with independently prepared inflorescence materials.

### RNA Extraction and Gene Expression Analysis

Total RNA was isolated after tissue collection above using TransZol reagent (TransGen Biotech); genomic DNA contamination was eliminated with DNase I (Roche). M-MLV reverse transcriptase (Thermo Scientific) was used for first-strand cDNA synthesis. qPCR was conducted in technical triplicates on a Bio-Rad CFX Connect real-time PCR system using SYBR Green PCR master mix (DBI Bioscience). *UBQ* was used as the reference gene. Three to four biological replicates were performed; the results were analyzed with SPSS statistics 17.0 (IBM).

### RNA-seq data analysis

The RNA-seq data in *35Spro: AP1-GR ap1 cal* was collected from a published article (Chen et al., 2018). Heat map visualizes the expression patterns of DEGs based on FPMK (fragments per kilobase of transcript per million mapped reads). The resulting p-values were adjusted for multiple comparisons by false discovery rate (FDR).

### *in Situ* Hybridization

For the *ARF3* mRNA probe, the *ARF3* coding region was amplified by RT-PCR and cloned into pGEM-T-easy (Promega). The resulting plasmid was digested with SpeI and transcribed with T7 RNA polymerase to generate the antisense probe. *In situ* hybridization was performed as previously described (Liu et al., 2011).

### ChIP Assay

ChIP was performed as previously described (Liu et al., 2011). Inflorescences were collected as above and ground in liquid nitrogen and crosslinked in 1% (w/v) formaldehyde (Sigma-Aldrich). Chromatin was extracted with M1, M2, M3 buffers and sonicated into DNA fragments of 500-1,000 bp. The lysate was precleared by incubation with 50 μL protein-A agarose beads (Roche) for 1 h and then incubated with anti-GFP (Abcam) antibodies overnight. The bound chromatin was purified on columns by using Qiagen Plasmid Extraction kit. qPCR was performed in technical triplicates.

### Histochemical Staining and Quantitative Analysis of β-Glucuronidase (GUS) Activity

For GUS staining, plant materials were immersed in GUS staining buffer (Leagene) and incubated for several hours in the dark at 37°C. Chlorophyll was removed by incubation in 70% (v/v) ethanol. Quantitative analysis of GUS activity was performed as described (Zhang et al., 2018).

### Accession Numbers

Sequence data from this article can be found in the Arabidopsis Genome Initiative or GenBank/EMBL data libraries under the following accession numbers: *AG* (At4g18960), *CRE*/*AHK4* (At2g01830), *ARF3* (At2g33860), *BOP1* (At3g57130), *BOP2* (At2g41370), *CUC1* (At3g15170), *CUC2* (At5g53950), *CUC3* (At1g76420), *eIF4A* (At3g13920), *MP*/*ARF5* (At1g19850), *STM* (At1g62360), *TEC3* (At2g28080), *UBQ* (At3g62250), and *WUS* (At2g17950).

## SUPPLEMENTAL INFORMATION

Supplemental information

## AUTHOR CONTRIBUTIONS

X.L., C.L. and K.Z. conceived and designed the project. K.Z., H.Z., Y.P. and L.G. performed the experiments with the help of S.T. J.W., Y.F. and C.W. conducted phenotypic statistics. T.L., Y.Z., H.S., Z.B. and J.D. analyzed data. K.Z. and P.Q. performed the confocal imaging. K.Z. and X.L. wrote the manuscript.

## ACKNOWLEDGMENTS

We thank all members of the Liu Lab for helpful discussion. This work was supported by grants from the National Science Foundation of China (31900623 to K.Z.; 31970824 to X.L.); the Natural Science Foundation of Hebei Province in China (C2019204041; C2021205013); the project (2020HBQZYC004; A202105008) from Hebei province; the Science and Technology Research Project of University of Hebei Province, China (QN202037). The study was funded by State Key Laboratory of North China Crop Improvement and Regulation (NCCIR2021ZZ-7).

---

**Supplemental Figure 1. L*er* and *arf3-29* mature plants**.

*arf3-29* mutants produce more siliques with delayed GPA than L*er*. Bars = 1 cm.

**Supplemental Figure 2. Distribution of *ARF3* mRNA and protein in IM and early stages of FMs**.

**A–D**, *ARF3* expression in IM (A) and FM at stage 3 (B), stage 6 (C) and stage 9–10

(D) examined by *in situ* hybridization. Bars = 50 μm.

**Supplemental Figure 3. The nls restricts the subcellular localization of ARF3**.

**A and B**, Subcellular localization of ARF3-GFP (A) and ARF3-nls-GFP (B). Bars = 5 μm.

**Supplemental Figure 4. Cell number in the SAM L1 layer of the indicated genotypes**.

**A–D**, Longitudinal section view of the indicated genotypes. **E**, Cell number of the SAM L1 layer of the plants in A-D. \*\**P* < 0.01 (Student’s t test). Bars = 25 μm in A– D.

**Supplemental Figure 5. Percentages of abnormal stamens in the indicated genotypes**.

**Supplemental Figure 6. Schematic diagram of the genomic regions of ARF3 target genes**.

Arrows indicate the transcription start site (+1). Dark gray rectangles, untranslated regions; black rectangles, exons; black lines, introns; red lines, fragments examined by ChIP-qPCR; green rectangles, ChIP-seq peaks regions (Simonini et al, 2017). Bar = 500 bp.

**Supplemental Figure 7. Representative siliques and phenotypes of the indicated genotypes**.

**A**–**D**, Representative siliques and plants of *ag-10* (A), *arf3-29 ag-10* (B), *arf3-29 ag-10 ARF3pro:ARF3-GFP* (C) and *arf3-29 ag-10 ARF3pro:ARF3-nls-GFP* (D). **E– H**, FM determinacy in the indicated genotypes: *ag-10* (E), *ag-10 arf3-29* (F), *ag-10 arf3-29 ARF3pro:ARF3-GFP* (G), *ARF3pro:ARF3-nls-GFP* (H). White arrow, hyperplastic tissue; yellow arrow, seed. **I**, Schematic diagrams of silique phenotypes of the indicated genotypes. Each column represents a single plant, and each square represents a silique. Green, normal siliques; yellow, intermediate-type siliques (silique bear seeds containing hyperplastic tissue inside); magenta, severe indeterminacy-type siliques (silique containing hyperplastic tissue inside without seeds). **J**, Percentages of total siliques from each type. Bars = 1 cm in A–D and 1 mm in E–G.

**Supplemental Figure 8. Expression of *AHK4pro:GUS***.

**A and B**, Expression of *AHK4pro:GUS* in L*er* (A) and *arf3-29* (B) inflorescences under the same staining conditions. Bars = 25 μm.

**Supplemental Figure 9. Representative size of SAM in the indicated genotypes. A–D**, Representative SAM of *wus-7* (A), *wus-7 arf3-29* (B), *cre1-10* (C) and *cre1-10 arf3-29* (D). **E**, Boxplot representation of SAM size distribution of the indicated genotypes. Statistical test: Student’ t test (p); Effect size: Hedges’ coefficient (g). Bars = 25 μm in A–D.

**Supplemental Table 1. Floral organ numbers in L*er* and *arf3-29* plants.**

**Supplemental Data 1. Expression patterns of genes in auxin signaling during early floral development in *ap1 cal 35Spro:AP1-GR***.

**Supplemental Data 2. List of primers used in this study**.

## REFERENCES

Bartrina, I., Otto, E., Strnad, M., Werner, T., and Schmulling, T. (2011). Cytokinin regulates the activity of reproductive meristems, flower organ size, ovule formation, and thus seed yield in Arabidopsis thaliana. Plant Cell 23, 69–80.

Bencivenga, S., Serrano-Mislata, A., Bush, M., Fox, S., and Sablowski, R. (2016). Control of Oriented Tissue Growth through Repression of Organ Boundary Genes Promotes Stem Morphogenesis. Dev Cell 39, 198–208.

Bilsborough, G.D., Runions, A., Barkoulas, M., Jenkins, H.W., Hasson, A., Galinha, C., Laufs, P., Hay, A., Prusinkiewicz, P., and Tsiantis, M. (2011). Model for the regulation of Arabidopsis thaliana leaf margin development. Proceedings of the National Academy of Sciences of the United States of America 108, 3424–3429.

Brand, U., Fletcher, J.C., Hobe, M., Meyerowitz, E.M., and Simon, R. (2000). Dependence of stem cell fate in Arabidopsis on a feedback loop regulated by CLV3 activity. Science (New York, NY) 289, 617–619.

Byrne, M.E., Groover, A.T., Fontana, J.R., and Martienssen, R.A. (2003). Phyllotactic pattern and stem cell fate are determined by the Arabidopsis homeobox gene BELLRINGER. Development 130, 3941–3950.

Cao, X., He, Z., Guo, L., and Liu, X. (2015). Epigenetic Mechanisms Are Critical for the Regulation of WUSCHEL Expression in Floral Meristems. Plant physiology 168, 1189–1196.

Carles, C.C., and Fletcher, J.C. (2003). Shoot apical meristem maintenance: the art of a dynamic balance. Trends in plant science 8, 394–401.

Chandler, J.W. (2012). Floral meristem initiation and emergence in plants. Cell Mol Life Sci 69, 3807–3818.

Chang, W., Guo, Y., Zhang, H., Liu, X., and Guo, L. (2020). Same Actor in Different Stages: Genes in Shoot Apical Meristem Maintenance and Floral Meristem Determinacy in Arabidopsis. Frontiers in Ecology and Evolution 8, 89.

Chen, D., Yan, W., Fu, L.Y., and Kaufmann, K. (2018). Architecture of gene regulatory networks controlling flower development in Arabidopsis thaliana. Nature communications 9, 4534.

Cheng, Z.J., Wang, L., Sun, W., Zhang, Y., Zhou, C., Su, Y.H., Li, W., Sun, T.T., Zhao, X.Y., Li, X.G., et al. (2013). Pattern of auxin and cytokinin responses for shoot meristem induction results from the regulation of cytokinin biosynthesis by AUXIN RESPONSE FACTOR3. Plant physiology 161, 240–251.

Chickarmane, V.S., Gordon, S.P., Tarr, P.T., Heisler, M.G., and Meyerowitz, E.M. (2012). Cytokinin signaling as a positional cue for patterning the apical-basal axis of the growing Arabidopsis shoot meristem. Proc Natl Acad Sci U S A 109, 4002–4007.

Chung, Y., Zhu, Y., Wu, M.F., Simonini, S., Kuhn, A., Armenta-Medina, A., Jin, R., Ostergaard, L., Gillmor, C.S., and Wagner, D. (2019). Auxin Response Factors promote organogenesis by chromatin-mediated repression of the pluripotency gene SHOOTMERISTEMLESS. Nature communications 10, 886.

Clough, S.J., and Bent, A.F. (1998). Floral dip: a simplified method for Agrobacterium-mediated transformation of Arabidopsis thaliana. The Plant journal 16, 735–743.

Daum, G., Medzihradszky, A., Suzaki, T., and Lohmann, J.U. (2014). A mechanistic framework for noncell autonomous stem cell induction in Arabidopsis. Proc Natl Acad Sci U S A 111, 14619–14624.

Gaillochet, C., Daum, G., and Lohmann, J.U. (2015). O cell, where art thou? The mechanisms of shoot meristem patterning. Current opinion in plant biology 23, 91–97.

Gaillochet, C., and Lohmann, J.U. (2015). The never-ending story: from pluripotency to plant developmental plasticity. Development 142, 2237–2249.

Gaillochet, C., Stiehl, T., Wenzl, C., Ripoll, J.J., Bailey-Steinitz, L.J., Li, L., Pfeiffer, A., Miotk, A., Hakenjos, J.P., Forner, J., et al. (2017). Control of plant cell fate transitions by transcriptional and hormonal signals. Elife 6.

Gordon, S.P., Chickarmane, V.S., Ohno, C., and Meyerowitz, E.M. (2009). Multiple feedback loops through cytokinin signaling control stem cell number within the Arabidopsis shoot meristem. Proc Natl Acad Sci U S A 106, 16529–16534.

Guo, L., Cao, X., Liu, Y., Li, J., Li, Y., Li, D., Zhang, K., Gao, C., Dong, A., and Liu, X. (2018). A chromatin loop represses WUSCHEL expression in Arabidopsis. The Plant journal: for cell and molecular biology 94, 1083–1097.

Higuchi, M., Pischke, M.S., Mahonen, A.P., Miyawaki, K., Hashimoto, Y., Seki, M., Kobayashi, M., Shinozaki, K., Kato, T., Tabata, S., et al. (2004). In planta functions of the Arabidopsis cytokinin receptor family. Proceedings of the National Academy of Sciences of the United States of America 101, 8821–8826.

Jasinski, S., Piazza, P., Craft, J., Hay, A., Woolley, L., Rieu, I., Phillips, A., Hedden, P., and Tsiantis, M. (2005). KNOX action in Arabidopsis is mediated by coordinate regulation of cytokinin and gibberellin activities. Current biology: CB 15, 1560–1565.

Kurata, T., Okada, K., and Wada, T. (2005). Intercellular movement of transcription factors. Current opinion in plant biology 8, 600–605.

Lee, Z.H., Hirakawa, T., Yamaguchi, N., and Ito, T. (2019). The Roles of Plant Hormones and Their Interactions with Regulatory Genes in Determining Meristem Activity. International journal of molecular sciences 20.

Lin, T.F., Saiga, S., Abe, M., and Laux, T. (2016). OBE3 and WUS Interaction in Shoot Meristem Stem Cell Regulation. PLoS One 11, e0155657.

Liu, X., Dinh, T.T., Li, D., Shi, B., Li, Y., Cao, X., Guo, L., Pan, Y., Jiao, Y., and Chen, X. (2014). AUXIN RESPONSE FACTOR 3 integrates the functions of AGAMOUS and APETALA2 in floral meristem determinacy. The Plant journal: for cell and molecular biology 80, 629–641.

Liu, X., Kim, Y.J., Muller, R., Yumul, R.E., Liu, C., Pan, Y., Cao, X., Goodrich, J., and Chen, X. (2011). AGAMOUS terminates floral stem cell maintenance in Arabidopsis by directly repressing WUSCHEL through recruitment of Polycomb Group proteins. The Plant cell 23, 3654–3670.

Long, J.A., Moan, E.I., Medford, J.I., and Barton, M.K. (1996). A member of the KNOTTED class of homeodomain proteins encoded by the STM gene of Arabidopsis. Nature 379, 66–69.

Ma, Y., Miotk, A., Sutikovic, Z., Ermakova, O., Wenzl, C., Medzihradszky, A., Gaillochet, C., Forner, J., Utan, G., Brackmann, K., et al. (2019). WUSCHEL acts as an auxin response rheostat to maintain apical stem cells in Arabidopsis. Nature communications 10, 5093.

Matsubayashi, Y. (2003). Ligand-receptor pairs in plant peptide signaling. Journal of cell science 116, 3863–3870.

Miwa, H., Kinoshita, A., Fukuda, H., and Sawa, S. (2009). Plant meristems: CLAVATA3/ESR-related signaling in the shoot apical meristem and the root apical meristem. J Plant Res 122, 31–39.

Nemhauser, J.L., Feldman, L.J., and Zambryski, P.C. (2000). Auxin and ETTIN in Arabidopsis gynoecium morphogenesis. Development 127, 3877–3888.

Peaucelle, A., Morin, H., Traas, J., and Laufs, P. (2007). Plants expressing a miR164-resistant CUC2 gene reveal the importance of post-meristematic maintenance of phyllotaxy in Arabidopsis. Development 134, 1045–1050.

Pfeiffer, A., Wenzl, C., and Lohmann, J.U. (2017). Beyond flexibility: controlling stem cells in an ever changing environment. Current opinion in plant biology 35, 117–123.

Reddy, G.V., Heisler, M.G., Ehrhardt, D.W., Meyerowitz, E.M., Sakai, H., Hua, J., Chen, Q.G., Chang, C., Medrano, L.J., Bleecker, A.B., et al. (2004). Real-time lineage analysis reveals oriented cell divisions associated with morphogenesis at the shoot apex of Arabidopsis thaliana. Development 131, 4225–4237.

Reinhardt, D., Mandel, T., and Kuhlemeier, C. (2000). Auxin regulates the initiation and radial position of plant lateral organs. Plant Cell 12, 507–518.

Riou-Khamlichi, C., Huntley, R., Jacqmard, A., and Murray, J.A. (1999). Cytokinin activation of Arabidopsis cell division through a D-type cyclin. Science 283, 1541–1544.

Sablowski, R. (2009). Cytokinin and WUSCHEL tie the knot around plant stem cells. Proceedings of the National Academy of Sciences of the United States of America 106, 16016–16017.

Schaller, G.E., Bishopp, A., and Kieber, J.J. (2015). The Yin-Yang of Hormones: Cytokinin and Auxin Interactions in Plant Development. The Plant cell 27, 44–63.

Scofield, S., Murison, A., Jones, A., Fozard, J., Aida, M., Band, L.R., Bennett, M., and Murray, J.A.H. (2018). Coordination of meristem and boundary functions by transcription factors in the SHOOT MERISTEMLESS regulatory network. Development 145.

Sessions, A., Nemhauser, J.L., McColl, A., Roe, J.L., Feldmann, K.A., and Zambryski, P.C. (1997). ETTIN patterns the Arabidopsis floral meristem and reproductive organs. Development 124, 4481–4491.

Shi, B., Guo, X., Wang, Y., Xiong, Y., Wang, J., Hayashi, K.I., Lei, J., Zhang, L., and Jiao, Y. (2018). Feedback from Lateral Organs Controls Shoot Apical Meristem Growth by Modulating Auxin Transport. Dev Cell 44, 204–216 e206.

Simonini, S., Bencivenga, S., Trick, M., and Ostergaard, L. (2017). Auxin-Induced Modulation of ETTIN Activity Orchestrates Gene Expression in Arabidopsis. The Plant cell 29, 1864–1882.

Sun, B., and Ito, T. (2015). Regulation of floral stem cell termination in Arabidopsis. Frontiers in plant science 6, 17.

Sun, B., Xu, Y., Ng, K.H., and Ito, T. (2009). A timing mechanism for stem cell maintenance and differentiation in the Arabidopsis floral meristem. Genes & development 23, 1791–1804.

Tantikanjana, T., and Nasrallah, J.B. (2012). Non-cell-autonomous regulation of crucifer self-incompatibility by Auxin Response Factor ARF3. Proceedings of the National Academy of Sciences of the United States of America 109, 19468–19473.

Vernoux, T., Besnard, F., and Traas, J. (2010). Auxin at the shoot apical meristem. Cold Spring Harb Perspect Biol 2, a001487.

Wei, J., Qi, Y., Li, M., Li, R., Yan, M., Shen, H., Tian, L., Liu, Y., Tian, S., Liu, L., et al. (2020). Low-cost and efficient confocal imaging method for arabidopsis flower. Developmental biology 466, 73–76.

Yadav, R.K., Perales, M., Gruel, J., Girke, T., Jonsson, H., and Reddy, G.V. (2011). WUSCHEL protein movement mediates stem cell homeostasis in the Arabidopsis shoot apex. Genes & development 25, 2025–2030.

Zadnikova, P., and Simon, R. (2014). How boundaries control plant development. Curr Opin Plant Biol 17, 116–125.

Zhang, K., Wang, R., Zi, H., Li, Y., Cao, X., Li, D., Guo, L., Tong, J., Pan, Y., Jiao, Y., et al. (2018a). AUXIN RESPONSE FACTOR3 Regulates Floral Meristem Determinacy by Repressing Cytokinin Biosynthesis and Signaling. The Plant cell 30, 324–346.

Zhao, H., Liu, L., Mo, H., Qian, L., Cao, Y., Cui, S., Li, X., and Ma, L. (2013). The ATP-binding cassette transporter ABCB19 regulates postembryonic organ separation in Arabidopsis. PloS one 8, e60809.

Zhao, Z., Andersen, S.U., Ljung, K., Dolezal, K., Miotk, A., Schultheiss, S.J., and Lohmann, J.U. (2010). Hormonal control of the shoot stem-cell niche. Nature 465, 1089–1092.

Zürcher, E., Tavor-Deslex, D., Lituiev, D., Enkerli, K., Tarr, P.T., and Müller, B. (2013). A Robust and Sensitive Synthetic Sensor to Monitor the Transcriptional Output of the Cytokinin Signaling Network in Planta. Plant physiology 161, 1066–1075.

